# Cerebello-Basal Ganglia Networks and Cortical Network Global Efficiency

**DOI:** 10.1101/2021.04.05.438208

**Authors:** T. Bryan Jackson, Jessica A. Bernard

## Abstract

The cerebellum (CB) and basal ganglia (BG) each have topographically distinct functional subregions that are functionally and anatomically interconnected with cortical regions through discrete thalamic loops and with each other via disynaptic connections, with previous work detailing high levels of functional connectivity between these phylogenetically ancient regions. It was posited that this CB-BG network provides support for cortical systems processing, spanning cognitive, emotional, and motor domains, implying subcortical network measures are strongly related to cortical network measures (Bostan & Strick, 2018); however, it is currently unknown how network measures within distinct CB-BG networks relate to cortical network measures. Here, 122 regions of interest comprising cognitive and motor CB-BG networks and 7 canonical cortical resting-state were used to investigate whether the integration (quantified using global efficiency, GE) of cognitive CB-BG network (CCBN) nodes and their segregation from motor CB-BG network (MCBN) nodes is related to cortical network GE and segregation in 233 non-related, right-handed participants (Human Connectome Project-1200). CCBN GE positively correlated with GE in the default mode, motor, and auditory networks and MCBN GE positively correlated with GE in all networks except the default mode and emotional. MCBN segregation was related to MN segregation. These findings highlight the CB-BG network’s role in executive function, task switching, and verbal working memory. This work has implications for understanding cortical network organization and cortical-subcortical interactions in healthy adults and may help in deciphering subcortical differences seen in disease states.

---

Until recently, the cerebellum (CB), basal ganglia (BG), and their networks have been relatively understudied outside of the realms of motor behavior and learning. However, the CB, BG, and their associated computations support a wide array of animal behaviors that are associated with cortical function, including human verbal, emotional, and cognitive abilities (Chen & Desmond, 2005; Guell et al., 2018; for reviews, see Ward et al., 2013 and Schmahmann, 2019). Thought to independently modulate cortical processes, the CB is associated with prediction and error-correcting processes (reviewed in Wolpert et al., 2011), while the BG is typically associated with reward and reinforcement-learning processes (reviewed in Lee, Seo, & Jung, 2012). Non-human primate studies have detailed the separate, closed-loop, trans-thalamic pathways required for this long-range communication connecting the CB and BG with the cortex and, indirectly, with each other (Kelly & Strick, 2003, 2004). However, more recent studies have revealed a direct, subcortical connection between the CB and the BG (Bostan et al., 2010; Hoshi et al., 2005). Here, we provide evidence of a highly integrated and distinct CB-BG functional networks and investigate their role in scaffolding cortical networks.

The CB and BG are phylogenetically ancient (Bell, 2002; Grillner & Robertson, 2016; Herrick M.D., 1924; Smeets et al., 2000; Stephenson-Jones et al., 2012) and throughout mammalian evolution, the CB scaled in size with the neocortex, though this increase in volume was not uniform across the structure. CB volume in great apes and humans is increased relative to neocortical volume (Barton & Venditti, 2014), with posterior CB (Crus I and Crus II) volume showing the most expansion (Balsters et al., 2010). Importantly, the posterior CB has dense projections to prefrontal regions (Ramnani et al., 2006) and has been associated with cognition in humans (Argyropoulos et al., 2020; Balsters et al., 2013; Balsters & Ranganath, 2008; King et al., 2019; Schmahmann, 2019; Stoodley & Limperopoulos, 2016). Likewise, primate BG regions are volumetrically larger in relation to the neocortex when compared to insectivores and other earlier mammals (Stephan & Andy, 1964). This is particularly evident in the striatum, which is known to be anatomically and functionally linked with regions of the cortex associated with higher-order processing (i.e. dorsolateral prefrontal cortex) in humans (Di Martino et al., 2008; Draganski et al., 2008). With these expansions in mind, cortically-associated cognitive processing seems to be linked to CB and BG evolution (Bostan & Strick, 2018) likely via concurrent cortical expansion; however, direct investigations of the relationships between CB-BG network and cortical network interactions within healthy subjects remain rare. Specifically, Bostan and Strick (2018) have suggested that the CB and BG may provide a foundation for the evolution of these cortical networks.

The discovery of anatomical bidirectional subcortical circuits in non-human primates provides a method of communication for and supports the concept of a CB-BG network (Bostan et al., 2010; Hoshi et al., 2005). Evidence from *in vivo* human studies parallel these findings. For example, motor-associated CB and BG nodes (e.g. Lob V and Lob VI of the CB; dorsal caudal and rostral putamen of the BG) are functionally connected with one another at rest; while cognition-associated regions of the CB and BG (e.g. Crus I, Crus II, and Lob VI of the CB; inferior and superior ventral striatum, dorsal caudate, and ventral rostral putamen of the BG) are likewise functionally connected with one another at rest (Hausman, Jackson, Goen, & Bernard, 2019). Diffusion tractography (Pelzer et al., 2013), viral tract tracing (Bostan et al., 2010), and human functional connectivity (Bernard et al., 2012, 2013) studies suggest that CB-BG communication does not only exclusively rely on indirect communication via the separate, bi-directional trans- thalamic loops (Kelly & Strick, 2003, 2004). Instead, perhaps the CB and BG are part of an anatomically and functionally integrated network that evolved to support cortical processes, including cognition in humans (Bostan & Strick, 2018).

Insights from across both neurological and psychiatric disease states support the potential importance of effective CB-BG integration. Many BG-related disorders display differences in the CB and its networks, including Parkinson’s disease (Caligiore et al., 2016; DeLong & Wichmann, 2010; Wu & Hallett, 2013), dystonia (Argyelan et al., 2009; Neychev et al., 2008), and resting tremors (e.g., Asanuma, 2006). Those with schizophrenia show differences in both the CB and its networks (Anticevic et al., 2014; Ji et al., 2019; Mavroudis et al., 2017) and dopaminergic circuits within the BG (Mehler-Wex et al., 2006), and hypokinetic movements were associated with changes in CB-motor cortex connectivity (Walther et al., 2017). Even in healthy aging, there are differences in the CB-BG circuit (Hausman et al., 2019) and within the closed-loop cortico-thalamo circuits of both the CB and BG (Bernard et al., 2013; Bo et al., 2014; Griffanti et al., 2018). These findings suggest that the CB-BG network and cortical loops are important to normal function and alterations may contribute to psychiatric disorders and dysfunction.

Given the phylogenetic age of the CB and BG and their involvement in multiple cortical networks and disease states, it has been posited that these regions may provide scaffolding for cortical networks (Bostan & Strick, 2018). fMRI provides a useful way to look at intrinsic brain organization and has implications for understanding function (Karahanoğlu & Van De Ville, 2017). Resting-state fMRI was used to determine networks of functionally-connected regions that largely overlap with task positive networks that are engaged during intentional behavior (e.g. cingulo-opercular network – CON; fronto-parietal network - FPN; Fair et al., 2009) or during self-directed thought and internal mentation (i.e. the default mode network, DMN; Buckner et al., 2008; Fair et al., 2009; Fox et al., 2015). We posit that if cortical networks are supported by CB-BG interactions, then how easily information travels within a CB-BG network (the degree of integration/interconnectedness) may be associated with information travel within cortical functional networks.

Graph theoretical approaches are useful in quantifying connectivity patterns and thus determining a given network. To quantify network integration, we calculated network-level global efficiency (GE) for both a cognitive and a motor CB-BG network and 7 canonical resting-state networks. GE is a graph theory metric defined as the average inverse shortest path length and was designed to capture the efficiency of information transmission within a given network (Rubinov & Sporns, 2010; Sporns et al., 2004; Whitfield-Gabrieli & Nieto-Castanon, 2012). It is currently unknown how CB-BG GE relates to cortical network GE; however, we hypothesized that the CB and BG each provide a foundation for cortical processing, consistent with the assertions of Bostan and Strick (2018), and thus predicted that CB-BG resting-state network-level GE would correlate positively with the GE of domain-specific cortical networks. Specifically, we predicted that cognitive CB-BG network (CCBN) GE would positively correlate with cognitive network (FPN, CON, and DMN) GE, while motor CB-BG network (MCBN) GE would be positively correlated with motor (MN) and sensory network (visual – VN, auditory – AN) GE. Previous work associated CB Lob V and Lob VI with emotional processes (Riedel et al., 2015). We further predicted a positive correlation between the MCBN GE and emotional network (EN) GE. We were also interested in network segregation (Chan et al., 2014). Segregation as a measure quantifies how disconnected nodes are from the global network. Given the hypotheses that CB-BG networks support cortical networks (Bostan & Strick, 2018), we predicted that CCBN segregation would correlate positively with FPN, CON, and DMN segregation and MCBN segregation would correlate positively with VN, AN, EN and MN segregation.

## Method

### Participants – HCP

Preprocessed resting-state functional MRI (rsfMRI) data from the S1200 Release of Human Connectome Project (WU-UMN HCP Consortium) were downloaded for analysis. Only data for unrelated, right-handed (Handedness>24) individuals without psychiatric diagnoses were used. Data were considered for this study only if the participant attained a high school degree (SSAGA_Educ>11), reported no family history of mental illness (FamHist_*_None = 1), did not meet the DSM4 criteria for Alcohol Abuse or Dependence (SSAGA_Alc_D4_Ab_Dx != 5; SSAGA_Alc_D4_Dp_Dx != 5), and did not meet the DSM criteria for Marijuana Dependence (SSAGA_Mj_Ab_Dep = 0). To account for any potential familial similarities in brain structure and function, only one participant from each family unit was chosen. Using Excel’s Rand() function, each participant was assigned a random number and the participant with the largest number in their family was selected for inclusion in the analyses. Only participants with a complete set of data (one T1w structural and four resting-state fMRI sessions with the associated movement parameters) were used for analysis. These exclusions resulted in a sample size of 233 individuals ranging in age from 22 to 37 years (111 males, 122 females; see Supplemental Text 1). Detailed descriptions of each variable used to eliminate participants are available here: https://wiki.humanconnectome.org (see HCP Data Dictionary Public- Updated for the 1200 Subject Release).

### HCP image acquisition and preprocessing

rsfMRI data for each participant consisted of four 15-minute runs collected over 2 sessions on different days (1200 volumes, 720 ms TR, 2 mm isotropic voxels; see Van Essen et al. (2012) for full details on fMRI acquisition). Notably, there were no differences in functional connectivity between the two sessions (see Supplemental Text 2 for detailed analysis methods and results). Preprocessed T1w structural, functional, and motion regressor data were downloaded from the HCP S1200 Release Amazon Web Services repository. Details of the preprocessing pipeline for functional images are fully described in Glasser et al. (2013). Structural scans had undergone gradient distortion correction, bias field correction, and registration to the 0.8 mm resolution MNI brain using Multimodal Surface Matching (Glasser et al., 2013, 2016). Additionally, structural images underwent tissue type segmentation and functional images underwent smoothing (5 mm FWHM) and artifact detection (global signal z-value threshold: 5, subject motion threshold: 0.9 mm) using the CONN toolbox (v. 20b; Whitfield-Gabrieli & Nieto-Castanon, 2012), a Matlab-based application designed for functional connectivity analysis and MATALB R2019b on centOS 7. Functional data were then denoised using confound regressors for 5 temporal components each from the segmented CSF and white matter, 24 motion realignment parameters, signal and/or motion outliers, and the 1st order derivative from the effect of rest. Finally, data underwent linear detrending and bandpass filtering (0.008 - 0.09 Hz).

### Regions of interest

Using fslmaths, 3.5 mm spherical, binarized regions of interest (ROIs) representing two rsfMRI networks of CB and BG nodes (cognitive and motor CB-BG network), and 7 rsfMRI cortical networks (3 cognitive-associated, FPN, CON, DMN; 1 motor-associated, MN; 1 emotion-associated, EN; and two sensory-associated, VN, AN) were created with coordinates from previous works (FSL v.6.0.3: FMRIB’s Software Library, www.fmrib.ox.ac.uk/fsl; Jenkinson, Beckmann, Behrens, Woolrich, & Smith, 2012). CB-BG network ROIs were created using CB and BG coordinates previously used in our recent work (Hausman et al., 2019). Lob V and Lob VI of the CB and dorsal caudal and dorsal rostral putamen of the BG were used to define the MCBN; while, Crus I and Crus II of the CB and the inferior and superior ventral striatum, dorsal caudate, and ventral rostral putamen of the BG were used to define the CCBN. This division is in line with previous work parcellating each region based on resting-state and task-based functional connectivity and behavioral associations (Bernard et al., 2012; Di Martino et al., 2008; Salmi et al., 2010; Stoodley et al., 2012)

The cognitive networks were defined using ROIs from two task-positive rsfMRI networks, the cingulo-opercular (CON) and the fronto-parietal (FPN), and the default mode network (DMN), using coordinates originally reported by Fair et al. (2009). Motor network (MN) ROIs were created using coordinates based on Mayka, Corcos, Leurgans, & Vaillancourt (2006). Emotional Network (EN) ROIs were created using coordinates originally reported by Stein et al. (2007) and adapted using information from Deen, Pitskel, & Pelphrey, (2011). The visual and auditory sensory networks, VN and AN, were defined using coordinates reported by Cassady et al. (2019). Importantly, given the association the CB and BG have with cortical and limbic networks, especially the MN, we did not use any CB or BG ROIs in any of the cortical networks. This required change of the VN only, for which we removed an ROI for CB Lobule VI. Removing this ROI allowed us to directly correlate measures from within the CB-BG networks to measures from within cortical networks. To account for the use of both left and right ROIs in some cortical networks, all lateralized (defined as more than 3mm from the midline in the x direction) ROIs without a reported contralateral match were mirrored inthe x direction (see Supplemental Table 2 for parallel analysis using only the original ROIs). Supplemental Table 1 and Figure 1 provide details for the 122 resulting ROIs. ROIs will be referred to by their abbreviation in Table 1 from this point forward.

**Figure 1.**
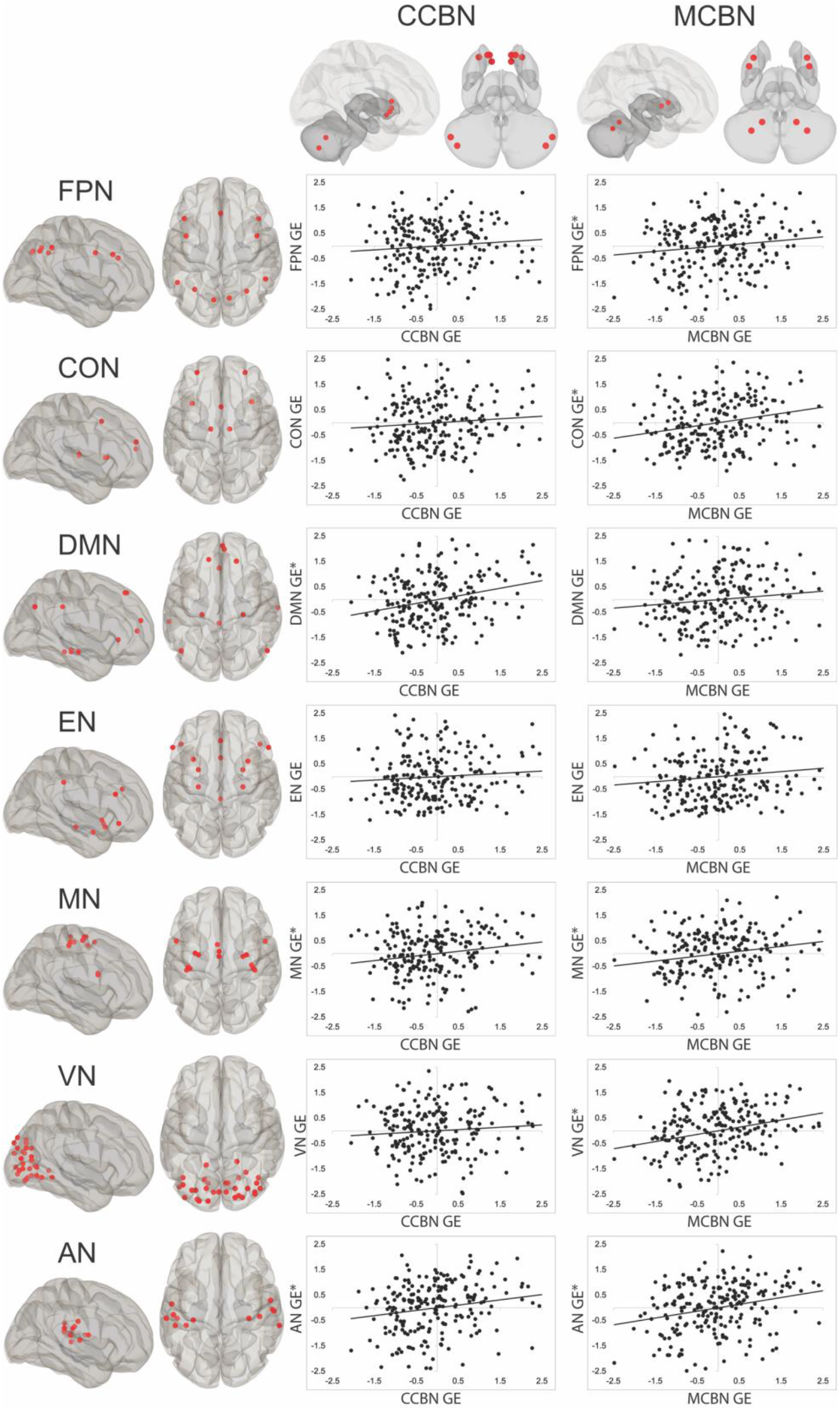
ROI locations for each network and scatterplots depicting the relationship between CCBM or MCBN GE and cortical network GE. To represent the partial correlations conducted in the analysis, standardized residuals of each CB-BG network (each controlling for the other) and standardized residuals of each cortical measure (controlling for the CB-BG network not in the analysis) were used. *Statistically significant correlation.

**Table 1.** β values and paired *t*-tests and for each significant ROI-ROI pair.

| ROI pairs | $\beta$ | $t$ (222) | $p$ (FDR) |
| --- | --- | --- | --- |
| R CrusI - R CrusII | 0.07 | 12.89 | 0.001 |
| R CrusI - R LobV | 0.01 | 2.88 | .008 |
| R CrusI - R LobVI | 0.02 | 5.09 | 0.001 |
| R CrusI - L VS<sub>i</sub> | 0.01 | 2.8 | .008 |
| R CrusI - L VRP | 0.02 | 6.16 | 0.001 |
| R CrusI - L DC | 0.02 | 5.83 | 0.001 |
| R CrusII - R LobV | 0.01 | 2.94 | .006 |
| R CrusII - R LobVI | 0.01 | 4.21 | 0.001 |
| R CrusII - L VS<sub>s</sub> | 0.01 | 3 | .006 |
| R CrusII - L VS<sub>i</sub> | -0.01 | -2.2 | .04 |
| R CrusII - L VRP | -0.01 | -2.84 | .007 |
| R CrusII - L DC | 0.01 | 3.97 | 0.001 |
| R LobV - R LobVI | 0.02 | 7.15 | 0.001 |
| R LobV - L DRP | 0.01 | 3.35 | 0.003 |
| R LobV - L DCP | 0.02 | 5.69 | 0.001 |
| R LobVI - L VS<sub>s</sub> | 0.01 | 2.38 | .02 |
| R LobVI - L VS<sub>i</sub> | 0.01 | 2.23 | .03 |
| R LobVI - L VRP | 0.01 | 3.01 | .004 |
| R LobVI - L DRP | 0.02 | 5.32 | 0.001 |
| R LobVI - L DCP | 0.01 | 4.23 | 0.001 |
| R LobVI - L DC | 0.02 | 4.54 | 0.001 |
| L VRP - L VS<sub>s</sub> | 0.07 | 16.93 | 0.001 |
| L VRP - L VS<sub>i</sub> | 0.04 | 12.7 | 0.001 |
| L VRP - L DC | 0.07 | 18.02 | 0.001 |
| L DRP - L VS<sub>s</sub> | 0.03 | 7.95 | 0.001 |
| L DRP - L VS<sub>i</sub> | 0.01 | 4.18 | 0.001 |
| L DRP - L VRP | 0.11 | 24.52 | 0.001 |
| L DRP - L DC | 0.05 | 13.83 | 0.001 |
| L DCP - L VS<sub>s</sub> | 0.01 | 3.44 | 0.001 |
| L DCP - L VRP | 0.06 | 15.84 | 0.001 |
| L DCP - L DRP | 0.12 | 26.71 | 0.001 |
| L DCP - L DC | 0.03 | 8.07 | 0.001 |
| L DC - L VS<sub>s</sub> | 0.05 | 12.17 | 0.001 |
| L DC - L VS<sub>i</sub> | 0.02 | 7.47 | 0.001 |
| L VS<sub>i</sub> - L VS<sub>s</sub> | 0.03 | 10.01 | 0.001 |

### fMRI participant-level correlation matrices

For each participant, a timeseries for each of the 122 ROIs was extracted and cross-correlated using bivariate analyses, resulting in a correlation matrix for each participant. All participant-level analyses were performed in the CONN toolbox (v. 20b; Whitfield-Gabrieli & Nieto-Castanon, 2012).

### Subcortical network analysis – replication

Subcortical ROIs originally used in our recent investigation of CB-BG connectivity in young and older adults (Hausman et al. 2019) were used to replicate the findings in a larger sample, again computed using the CONN toolbox (v. 20b; Whitfield-Gabrieli & Nieto-Castanon, 2012). Bivariate correlations were calculated by cross correlating each timeseries of four lateralized CB ROIs (R CrusI, R CrusII, R LobV, and R LobVI) and six lateralized BG ROIs (L VSi, L VSs, L VRP, L DRP, L DC, L DCP). Fisher z-transformed correlation matrices were computed and thresholded at *p*(FDR) = .05.

### Graph theory and correlational analyses

Average network-level graph theoretical measures were computed separately for each of the seven networks within the CONN toolbox (Whitfield-Gabrieli & Nieto-Castanon, 2012). Edges were defined as thresholded positive correlation coefficients (*β* > .1) and nodes were defined as the ROIs in a given network (Fair et al., 2009). Given previous findings that show a high degree of integration within each network (Fair et al., 2009; Geerligs et al., 2015; Hausman et al., 2019; Stein et al., 2007), we focused on network-level GE, a measure of how efficiently information in a network travels, as the primary measure of each network’s integration for our analyses here (Achard & Bullmore, 2007; Latora & Marchiori, 2001; Sheffield et al., 2016; Whitfield-Gabrieli & Nieto-Castanon, 2012). To test whether the GE of the CB-BG networks were related to the GE of the FPN, CON, DMN, EN, MN, VN, or AN, partial Pearson correlations were performed between the each of the CB-BG networks and each of the other networks (i.e., CCBN GE compared to FPN GE) using IBM SPSS 25 (IBM Corporation Armonk, NY, 2017). We also calculated the following network-level graph theoretical measures and performed the same analyses in order to compare the findings in an exploratory manner: betweenness centrality (average of the number of times each node is part of a shortest-path between any two ROIs), cost/degree (average of the proportion/number of edges that are connected to a given node - for our purposes, cost and degree are redundant measures and are discussed as one), and average path length (average number of edges traversed from ROI-to-ROI). For a full mathematical description of each of these measures and of GE, please see Achard & Bullmore, 2007; Latora & Marchiori, 2001; Rubinov & Sporns, 2010; Sporns et al., 2004 and Whitfield-Gabrieli & Nieto-Castanon, 2012. Bonferroni corrections were applied to all exploratory correlations.

### Network segregation analysis

Participant-level Fisher z-transformed correlation matrices were used to calculate within-network and between-network connectivity separately for each network (Chan et al., 2014). Within-network connectivity was defined as the average z-score for all ROI-to-ROI connections in a network. Between-network connectivity was defined differently for the CB-BG and cortical networks. For each of the CB-BG networks, between-network connectivity was defined as the average z-score of the ROI-to-ROI connections between nodes in the CCBN and nodes in the MCBN. For all cortical networks, between-network connectivity was defined as the average z-score of the ROI-to-ROI connections between all nodes in every other cortical network (leaving out all CB-BG nodes). This allowed us to test whether motor and cognitive node segregation within the CB and BG (and vice versa) was related to the segregation of cortical networks. Next, network segregation was calculated separately for each network using the following formula:

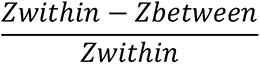

To answer the question of whether subcortical segregation predicted cortical segregation, Pearson correlations were performed between the CB-BG networks and each cortical network’s segregation score. We chose to use bivariate, as opposed to partial, correlations for the segregation analyses, due to the nature of how both CB-BG networks are measured when segregation is calculated.

## Results

### CB-BG connectivity network

Figure 2 depicts the connectivity patterns and Table 1 lists *t*-scores and beta values for each ROI-to-ROI pair in the lateralized CB-BG network analysis, performed to replicate our previous findings in a larger sample (Hausman et al., 2019). Broadly speaking, these results successfully replicate our past work showing robust positive connectivity between the CB and BG. Notably, these analyses did reveal 2 anticorrelations that were not seen in our prior work (Hausman et al., 2019). These anticorrelations were seen between R CrusII and both L VSi and L VRP. We suspect however, that the anticorrelations found here may be due to higher sensitivity and statistical power associated with the larger sample used in this investigation.

**Figure 2.**
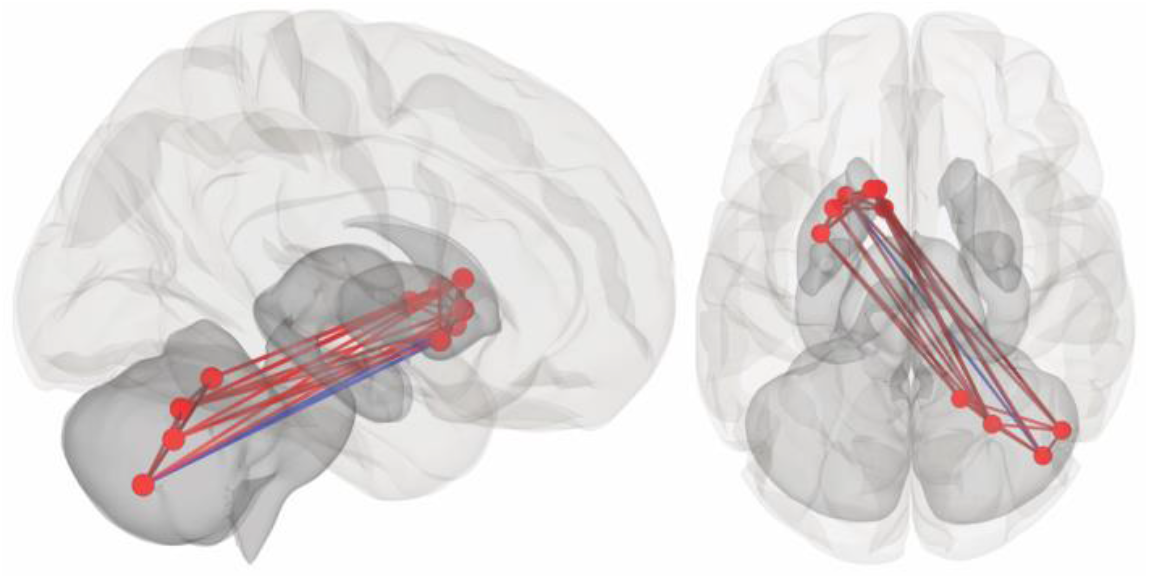
Right and superior views of connectivity patterns within right CB and left BG seeds.

### Subcortical-cortical graph theory network-level correlations

Table 2 lists partial Pearson correlation coefficients (*r*) and Figure 1 depicts the scatterplots for relationships between CB-BG and cortical network GE. A positive correlation was found between CCBN GE and DMN GE, in line with our predictions. We additionally found unexpected positive correlations between the GE of CCBN and that of MN and AN. No associations between CCBN GE and FPN and CON GE were found, contrary to our hypotheses. MCBN network GE positively correlated with the MN, VN, and AN, as predicted. Positive correlations were also found between MCBN GE and FPN and CON GE, unexpectedly. There were no associations between MCBN GE and EN GE. For a complete picture of cortical network integration, exploratory correlational analyses between cortical and limbic network GE are presented in Supplemental Table 3. Positive correlations were found between every network pairing except for FPN/EN.

**Table 2.**
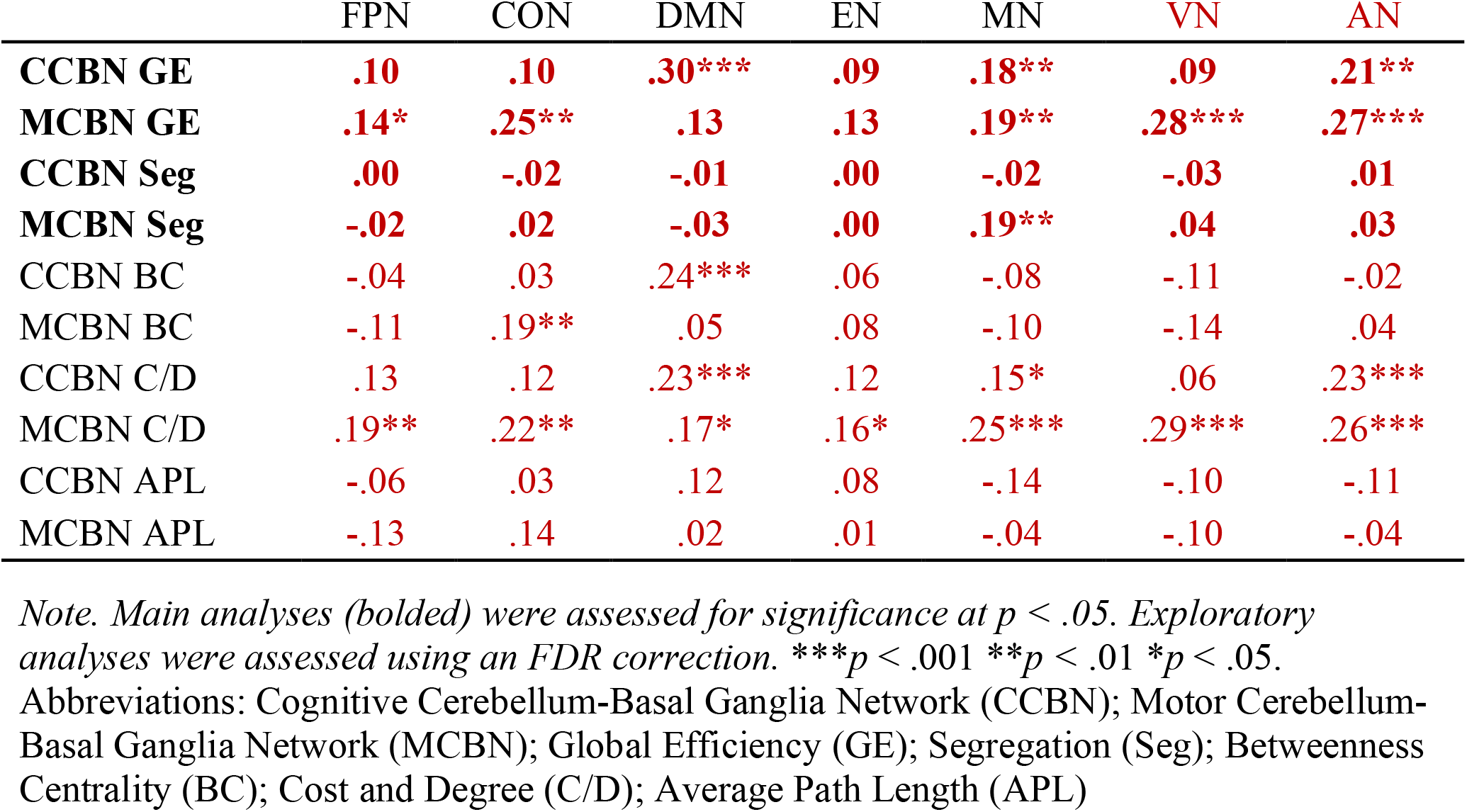
Pearson correlation values (*r’s)* for each subcortical-cortical pairing.

|  | FPN | CON | DMN | EN | MN | VN | AN |
| --- | --- | --- | --- | --- | --- | --- | --- |
| **CCBN GE** | **.10** | **.10** | **.30\*\*\*** | **.09** | **.18\*\*** | **.09** | **.21\*\*** |
| **MCBN GE** | **.14\*** | **.25\*\*** | **.13** | **.13** | **.19\*\*** | **.28\*\*\*** | **.27\*\*\*** |
| **CCBN Seg** | **.00** | **-.02** | **-.01** | **.00** | **-.02** | **-.03** | **.01** |
| **MCBN Seg** | **-.02** | **.02** | **-.03** | **.00** | **.19\*\*** | **.04** | **.03** |
| CCBN BC | -.04 | .03 | .24*** | .06 | -.08 | -.11 | -.02 |
| MCBN BC | -.11 | .19** | .05 | .08 | -.10 | -.14 | .04 |
| CCBN C/D | .13 | .12 | .23*** | .12 | .15* | .06 | .23*** |
| MCBN C/D | .19** | .22** | .17* | .16* | .25*** | .29*** | .26*** |
| CCBN APL | -.06 | .03 | .12 | .08 | -.14 | -.10 | -.11 |
| MCBN APL | -.13 | .14 | .02 | .01 | -.04 | -.10 | -.04 |

Note. Main analyses (bolded) were assessed for significance at $p < .05$ . Exploratory analyses were assessed using an FDR correction. \*\*\* $p < .001$ \*\* $p < .01$ \* $p < .05$ .
Abbreviations: Cognitive Cerebellum-Basal Ganglia Network (CCBN); Motor Cerebellum-Basal Ganglia Network (MCBN); Global Efficiency (GE); Segregation (Seg); Betweenness Centrality (BC); Cost and Degree (C/D); Average Path Length (APL)

Table 2 lists partial Pearson correlation coefficients (*r*) for relationships between CB-BG and cortical network segregation. CCBN segregation did not correlate with any cortical network segregation, contrary to our hypotheses. MCBN segregation correlated with MN, in line with our hypotheses. No other correlations involving segregation were found (*ps* > .06). We also explored correlations between cortical network segregation. We found significant positive correlations between CON and VN segregation (*r*(221) = .21, *p* = .01), and between MN and both sensory networks (VN, *r*(221) = .39, *p* < .001; AN, *r*(221) = .33, *p* < .001). We also found a significant negative correlation between FPN and AN segregation (*r*(221) = -.15, *p* = .04). No other significant correlations were found between any cortical network (*ps* > .07).

The exploratory correlations between CB-BG and cortical/limbic graph theory measures are also presented in Table 2. Cost and degree of the CCBN correlated with that of the DMN, MN, and AN, in line with the GE results. Cost and degree of the MCBN correlated with that of FPN, CON, MN, VN, and AN, mirroring the GE analyses, and additionally the DMN and EN. Betweenness centrality (BC) of the CCBN positively correlated with that of the DMN. MCBN network BC positively correlated with that of the CON. No other subcortical-cortical correlations were found when using betweenness centrality. No correlations were found using APL as a network metric.

## Discussion

Here, we performed novel correlational analyses relating cognitive and motor CB-BG network GE to cortical network GE to investigate whether the integration and separation of CB- BG networks correlates with that of cortical networks, as would be predicted given Bostan & Strick’s (2018) hypothesis that the CB-BG network is foundational to cortical networks and cognition. These results suggest some degree of specificity of cognitive/motor CB-BG networks as they relate to cortical networks. This has implications for understanding cortical network processing and cortical-subcortical interactions in healthy adults and may help in understanding the broad network-based changes seen in disease states.

First, we replicated our previous work (Hausman et al., 2019) investigating CB-BG connectivity at rest and report strong connectivity between these subcortical regions. Our findings are consistent with our prior work demonstrating robust connectivity between the CB and the BG, but this replication revealed several new findings as well. Within the CB, we found associations between every lateralized node, and a similar pattern again emerged within the BG. We also found that R LobVI is functionally connected to every BG ROI, reflecting R LobVI’s role in both motor and cognitive function (Balsters et al., 2013) and the functional connections to premotor, motor, and prefrontal area (Bernard et al., 2012). However, we also found anticorrelations for R CrusII - L VSi and R CrusII - L VRP connections. This may be indicative of delayed or lagged communication between these regions but the best interpretation of negative relationships between time courses is up for debate (Meszlényi et al., 2017).

Contrary to our predictions about task-positive cognitive networks, we found that FPN and CON GE were significantly related to the MCBN GE, yet we found only marginal associations between CCBN GE and FPN and CON GE. The relationship between FPN and MCBN integration may be due to the involvement of CB Lob VI in cognition and working memory, cognitive control, and response inhibition (Balsters et al., 2013; Bernard et al., 2020; Clark et al., 2020; Von der Gablentz et al., 2015), while the relationship with CON integration is perhaps due to the network’s association with alertness and error processing (Coste & Kleinschmidt, 2016; Neta et al., 2014). Regarding cognitive networks, it may be that efficiency within motor nodes of the CB-BG network may be more related to cognition than efficiency within the cognitive nodes; however, while there is a difference between the CB-BG networks in their relationships with the FPN, the effect sizes are similar (*rs* = .10 and .14), so caution must be taken in interpreting the differences.

In line with our predictions, the GE of the DMN, a task-negative network associated with self-reflective thought and mind-wandering (Fox et al., 2015), was related to CCBN GE, highlighting the role of the CB and BG in this key network. However, DMN segregation from cortical networks was not related to segregation of CCBN from the MCBN network, inconsistent with our predictions. It seems that while the integration of cognitive regions of the CB-BG network may support DMN processing, separation of CB-BG cognitive regions from motor systems is not related to DMN function in healthy adults.

The GE of both CB-BG networks correlated positively with MN GE, indicating that the CB and BG are involved not only with motor behavior, through its association with the MN (Bernard et al., 2014; Jackson et al., 2020; Kelly & Strick, 2003, 2004), but also with cognitive control via the prefrontal-cortical loops that innervate the CCBN (Bostan & Strick, 2018; Da Cunha et al., 2012; Haber, 2014; Schmidt et al., 2008; Sgambato-Faure et al., 2016; Yin et al., 2008). Additionally, MCBN segregation correlated positively with MN segregation, highlighting the importance of motor segregation in healthy young adults.

Sensory network GE positively correlated with MCBN GE, as expected due to previous functional parcellations of the CB and BG (Buckner et al., 2011; Di Martino et al., 2008) and their historical associations with motor behavior. Interestingly, CCBN GE also correlated with AN GE, possibly due to subvocalization and auditory loops used in cognition and verbal working memory (Rottschy et al., 2012). MCBN segregation correlated with AN segregation, indicating that, while CCBN integration may be important for subvocalization, the separation of motor from cognitive regions of the CB-BG network is important for sound and language processing, potentially due to CB-BG involvement in discriminating between subvocal cognition and sensory stimuli processing, as is thought to break down in psychosis-related disorders (e.g., Debbané et al., 2010).

No relationships were found between either the CCBN or MCBN and the EN. We originally predicted that due to the involvement of Lobule VI in emotional processes (Riedel et al., 2015), that MCBN GE and segregation would correlate positively with that of the EN. However, our null results indicate that efficiency and segregation of the cognitive and motor CB- BG networks is not related to the EN. We think this may be in part due to the method by which limbic regions within the EN communicate with the cortex. The amygdala and related structures project to cortical regions directly (MacLean, 1988), indicating limbic regions may have evolved to support cortical behavior independent of CB-BG interactions, perhaps to decrease the time it takes to relay important information like the identification of a threat.

To further understand the relationships between the CB-BG network and cortical networks, parallel exploratory analyses were conducted using other graph theory measures. Cost and degree analyses generally replicated the main GE analyses (see Table 2). This was expected, as these are measures of the number of connections in a network and are proxies for network efficiency (Rubinov & Sporns, 2010). BC, a measure of the average proportion of times that an ROI in a network is part of the shortest ROI-to-ROI path within the network (Rubinov & Sporns, 2010), showed a relationship between the CCBN and the DMN, in line with GE analyses. MCBN BC correlated with the CON and VN BC, highlighting motor involvement with alertness and attention (Coste & Kleinschmidt, 2016; Neta et al., 2014), partially mirroring the GE analyses. Similarly, APL only related the MCBN with the CON and VN, indicating that BC and APL are potentially not as sensitive as cost and degree or GE in determining relationships between subcortical-cortical pairs. This may be because not all within-network ROIs are connected, resulting in ROIs with zero efficiency being left out of the equations for C/D, APL, and BC and thus failing to fully capture the efficiency of the network. GE solves this issue by including ROI connections that have zero efficiency (Rubinov & Sporns, 2010), and provides further support for using GE in our analyses. Together, these exploratory findings help to confirm the idea that subcortical integration is related to the integration of large-scale cortical networks, though the degree to which this is the case varies by integration measure.

## Limitations

While this work provides important new insights into the interplay between subcortical CB- BG networks and those in the cortex, there are several limitations to consider. For each network, we used an unequal number of ROIs that represent a subset of the possible ROIs that could be used to investigate these networks. These choices were rooted in the literature of canonical resting-state networks (Deen et al., 2011; Fair et al., 2009; Mayka et al., 2006; Stein et al., 2007) but accounting for the disparate number of regions, using a whole-brain parcellation approach, or defining ROIs within a network using a more data-driven method may produce different results and these methods warrant their own studies. Future studies may also benefit from simply increasing the number of ROIs, allowing for higher resolution when capturing network measures. CB-BG networks may generally benefit from adding ROIs that incorporate additional areas within each region (i.e. including additional domain-associated CB nodes in each network; including BG regions other than the striatum, caudate, and putamen). In particular, a meta-analysis detailed emotional processing withing CB Lobules IV, V, VI, VIII, and IX (E et al., 2014) and perhaps a relationship with the EN would be found using different nodes or an additional CB-BG network. Relatedly, there are additional cortical networks that can be included. For example, we examined the CON network but did not distinguish between the ventral and dorsal attention systems. Additional ROIs and networks may help better capture information travel and integration and assist in deciphering complicated cortical-CB-BG relationships.

Additionally, although there are CB and BG nodes known to be involved in many of the cortical resting-state networks, the cortical networks chosen for use did not include any CB or BG ROIs (a single CB node within the VN was removed prior to analyses). This allowed us to directly compare CB-BG and cortical network patterns. While necessary for our purposes, this method inherently truncates the size of the functional networks to only cortical nodes, potentially limiting the accuracy of the network measure in the context of how these networks are recruited during task processing. Lastly, we used classic time-series autocorrelational analyses to build an ROI-to-ROI table. Other methods that can account for temporal lag like dynamic time warping (Meszlényi 2017) or sliding window analyses may reveal additional details about the temporal order of activation in within-network ROI relationships that are used to calculate the graph theory measures.

## Conclusions

These findings suggest CB-BG network integration is related to cortical network integration, and cognitive/motor segregation is important for cortical processing, in support of theories suggesting that the CB and BG each serve a foundational role for learning and cognition in support of domain-specific cortical networks (Bostan & Strick, 2018). We also showed that, while regions associated with similar behavioral domains (cognitive, motor, etc.) tend to be highly functionally connected (Bernard et al., 2013; Hausman et al., 2019), CB and BG subregions are generally highly functionally connected, even between regions associated with different behavioral domains and cortical circuitsExploring the CB-BG relationship and their cortical projections within the context of psychopathologies would be interesting and worthwhile. These findings detail network dynamics in a large representative sample and may help to elucidate network-related changes in clinical or older adult populations in the future.

## Acknowledgements

Portions of this research were conducted with the advanced computing resources provided by Texas A&M High Performance Research Computing.

## Supplemental Text

1. 100610, 101309, 102614, 102715, 102816, 103111, 103818, 104012, 104416, 105014, 105115, 105216, 105620, 105923, 106016, 106824, 107422, 108020, 108121, 108323, 108525, 109325, 109830, 111009, 111312, 111413, 111716, 112314, 112516, 112920, 113215, 113922, 114621, 114823, 114924, 115017, 116726, 117122, 118124, 118730, 118831, 119126, 119732, 119833, 120414, 120515, 121416, 122317, 122620, 123521, 123925, 124422, 125222, 126426, 126628, 127630, 127832, 128026, 128632, 128935, 129028, 129331, 129634, 130114, 130316, 130417, 130619, 130821, 131217, 132017, 132118, 133019, 133928, 134223, 134627, 135225, 136833, 137027, 137532, 138130, 138332, 138837, 140319, 143426, 144125, 144832, 145127, 146129, 146533, 149741, 149842, 150423, 151324, 151425, 151728, 151930, 152427, 153126, 153227, 153833, 154229, 154734, 156031, 156233, 156334, 156536, 157437, 157942, 159239, 159744, 161630, 162228, 162733, 163432, 164030, 164131, 164939, 167036, 167440, 168139, 169545, 172029, 172332, 172433, 172534, 173435, 173738, 173940, 174841, 175742, 176542, 177241, 178142, 178849, 178950, 179245, 180230, 180533, 180735, 180836, 182739, 182840, 185442, 185846, 186040, 188145, 189349, 189450, 190031, 191033, 192136, 195041, 195647, 196346, 198249, 198855, 199352, 202113, 204016, 204420, 206323, 206727, 208125, 211114, 211316, 212015, 212217, 213522, 214423, 219231, 220721, 236130, 245333, 275645, 281135, 285446, 286650, 287248, 299760, 304020, 309636, 310621, 311320, 314225, 316633, 321323, 329844, 336841, 349244, 350330, 358144, 361941, 381543, 385046, 391748, 397154, 397760, 414229, 419239, 459453, 468050, 510225, 516742, 529549, 529953, 547046, 552241, 561949, 567759, 571144, 580751, 589567, 618952, 622236, 675661, 680250, 680452, 748662, 757764, 789373, 835657, 849264, 97998

2. Data were collected on two different days. Each scanning session consisted of two 15-minute scans, resulting in four 15-minute scans total. The runs in each session were averaged together and the connectivity matrices for each session were contrasted against one another at a group-level using CONN. A false discovery rate correction was applied. No differences were found between the two sessions, indicating stable network connectivity patterns across sessions, even when data are collected on different days, *ps* > .9.

---

**Supplemental Table 1.**
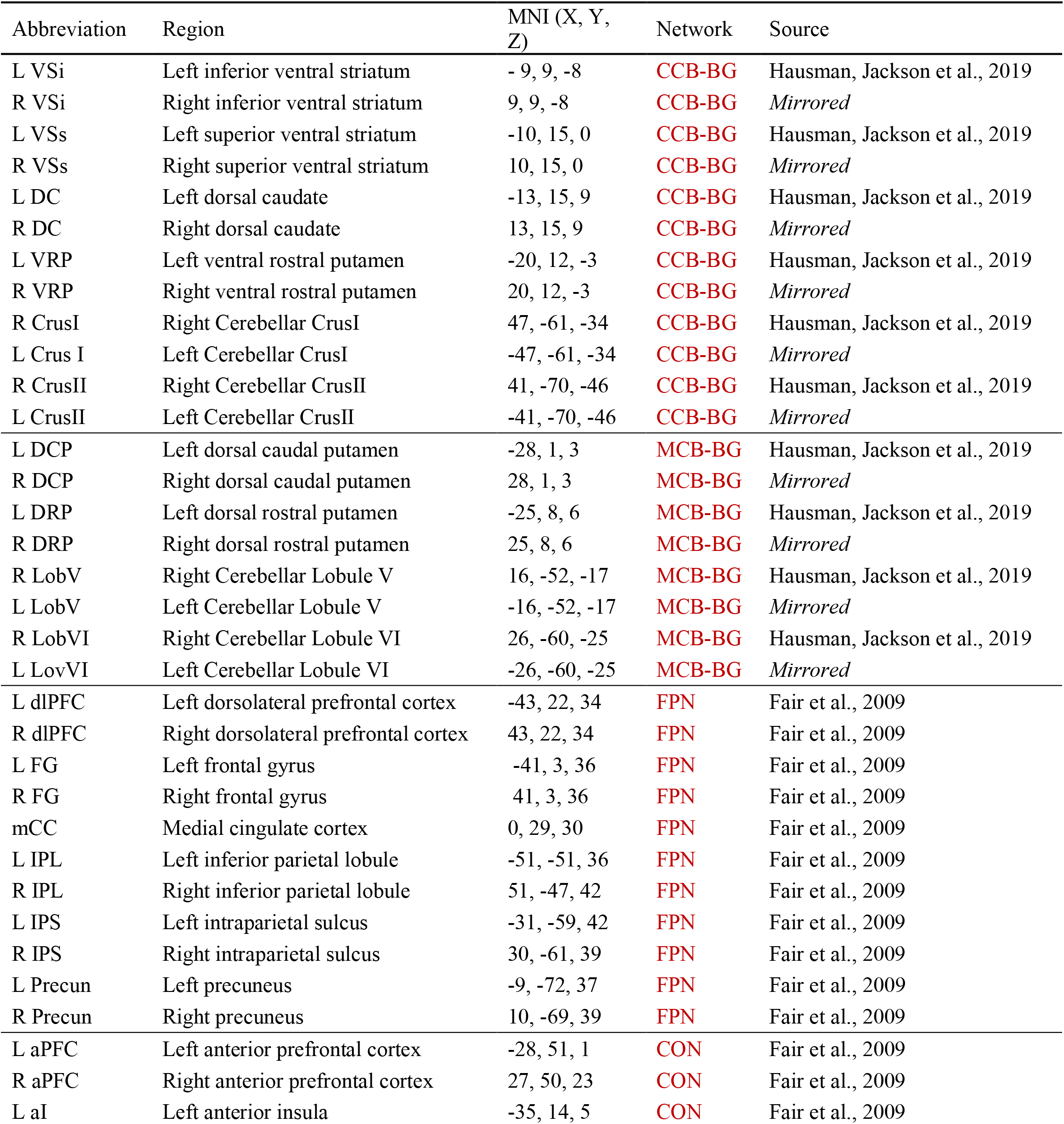
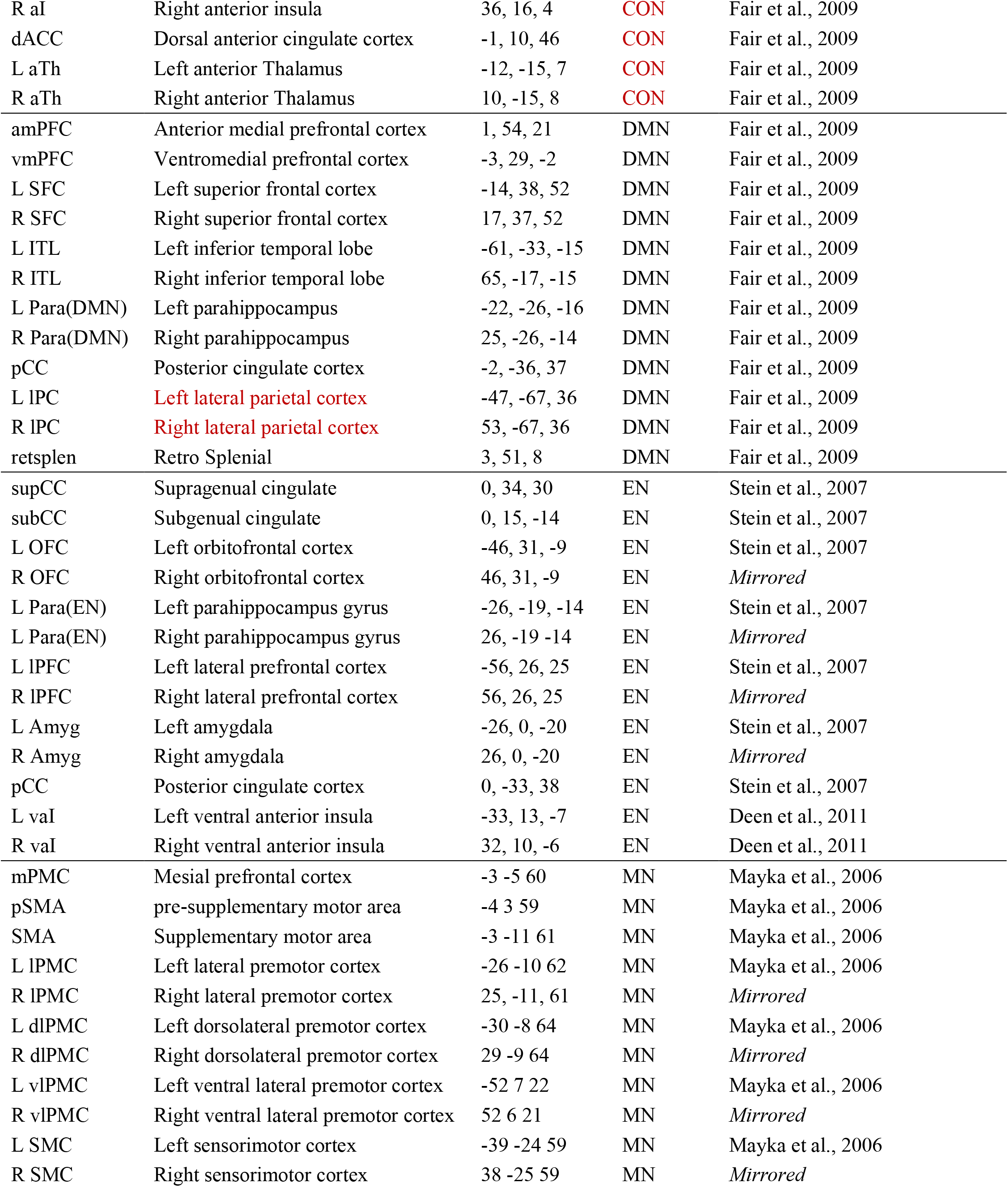
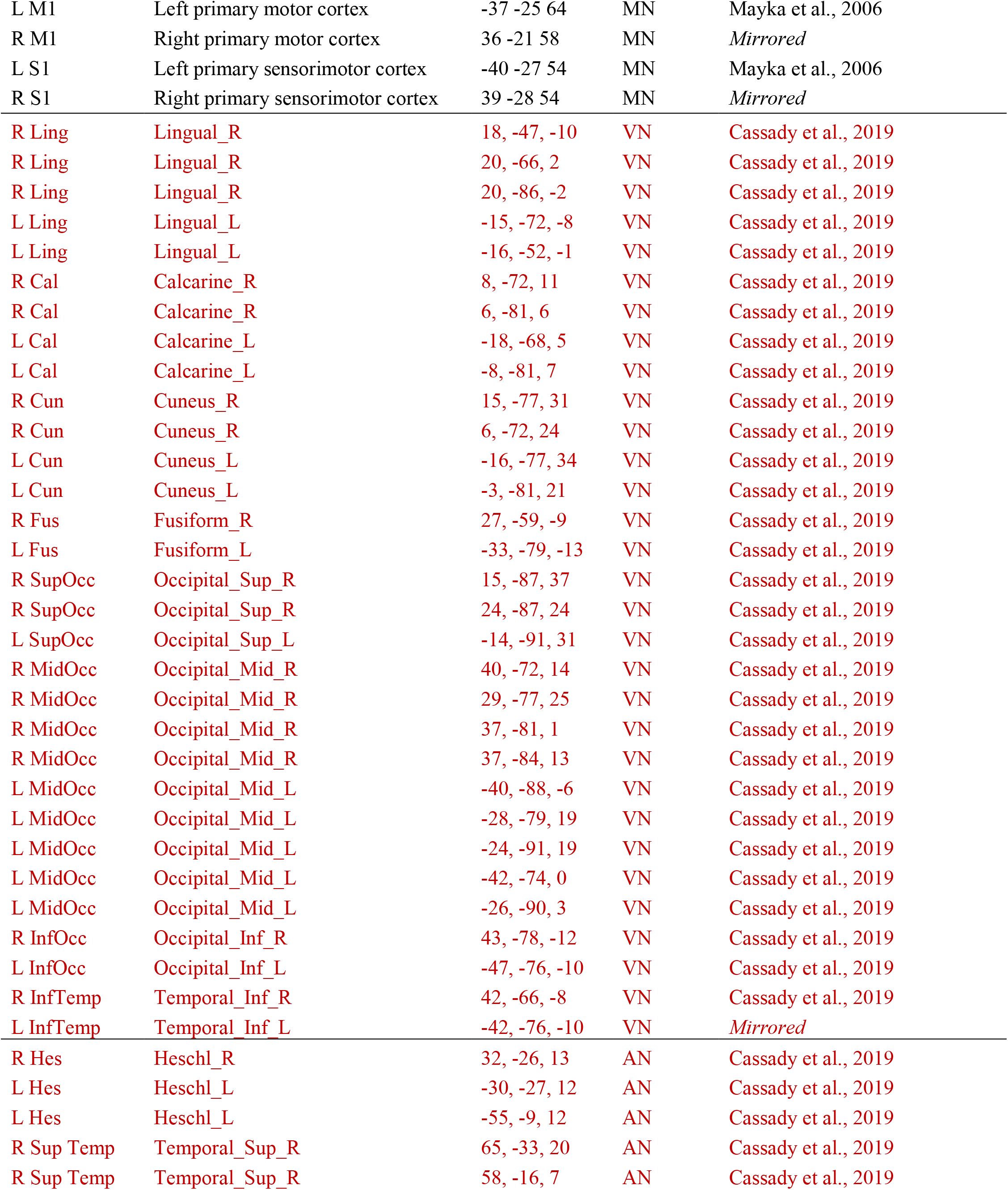
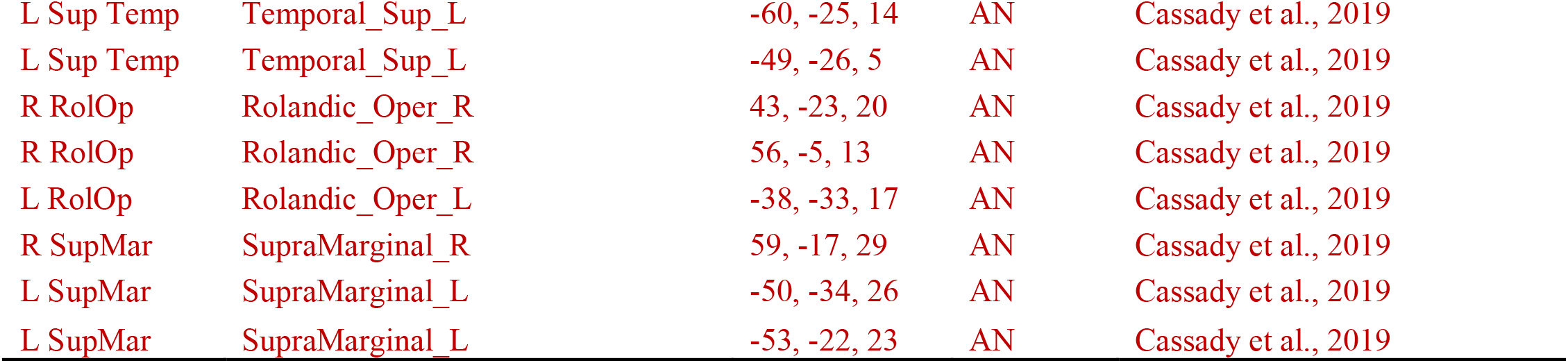
Abbreviations, region names, MNI coordinates, network association, and source for each ROIs used in the analyses. All ROIs are 3.5mm, spherical, and binarized. ROIs with the source “*Mirrored*” were created using its contralateral pair.

| Abbreviation | Region | MNI (X, Y, Z) | Network | Source |
| --- | --- | --- | --- | --- |
| L VSi | Left inferior ventral striatum | - 9, 9, -8 | CCB-BG | Hausman, Jackson et al., 2019 |
| R VSi | Right inferior ventral striatum | 9, 9, -8 | CCB-BG | *Mirrored* |
| L VSs | Left superior ventral striatum | -10, 15, 0 | CCB-BG | Hausman, Jackson et al., 2019 |
| R VSs | Right superior ventral striatum | 10, 15, 0 | CCB-BG | *Mirrored* |
| L DC | Left dorsal caudate | -13, 15, 9 | CCB-BG | Hausman, Jackson et al., 2019 |
| R DC | Right dorsal caudate | 13, 15, 9 | CCB-BG | *Mirrored* |
| L VRP | Left ventral rostral putamen | -20, 12, -3 | CCB-BG | Hausman, Jackson et al., 2019 |
| R VRP | Right ventral rostral putamen | 20, 12, -3 | CCB-BG | *Mirrored* |
| R CrusI | Right Cerebellar CrusI | 47, -61, -34 | CCB-BG | Hausman, Jackson et al., 2019 |
| L Crus I | Left Cerebellar CrusI | -47, -61, -34 | CCB-BG | *Mirrored* |
| R CrusII | Right Cerebellar CrusII | 41, -70, -46 | CCB-BG | Hausman, Jackson et al., 2019 |
| L CrusII | Left Cerebellar CrusII | -41, -70, -46 | CCB-BG | *Mirrored* |
| L DCP | Left dorsal caudal putamen | -28, 1, 3 | MCB-BG | Hausman, Jackson et al., 2019 |
| R DCP | Right dorsal caudal putamen | 28, 1, 3 | MCB-BG | *Mirrored* |
| L DRP | Left dorsal rostral putamen | -25, 8, 6 | MCB-BG | Hausman, Jackson et al., 2019 |
| R DRP | Right dorsal rostral putamen | 25, 8, 6 | MCB-BG | *Mirrored* |
| R LobV | Right Cerebellar Lobule V | 16, -52, -17 | MCB-BG | Hausman, Jackson et al., 2019 |
| L LobV | Left Cerebellar Lobule V | -16, -52, -17 | MCB-BG | *Mirrored* |
| R LobVI | Right Cerebellar Lobule VI | 26, -60, -25 | MCB-BG | Hausman, Jackson et al., 2019 |
| L LovVI | Left Cerebellar Lobule VI | -26, -60, -25 | MCB-BG | *Mirrored* |
| L dlPFC | Left dorsolateral prefrontal cortex | -43, 22, 34 | FPN | Fair et al., 2009 |
| R dlPFC | Right dorsolateral prefrontal cortex | 43, 22, 34 | FPN | Fair et al., 2009 |
| L FG | Left frontal gyrus | -41, 3, 36 | FPN | Fair et al., 2009 |
| R FG | Right frontal gyrus | 41, 3, 36 | FPN | Fair et al., 2009 |
| mCC | Medial cingulate cortex | 0, 29, 30 | FPN | Fair et al., 2009 |
| L IPL | Left inferior parietal lobule | -51, -51, 36 | FPN | Fair et al., 2009 |
| R IPL | Right inferior parietal lobule | 51, -47, 42 | FPN | Fair et al., 2009 |
| L IPS | Left intraparietal sulcus | -31, -59, 42 | FPN | Fair et al., 2009 |
| R IPS | Right intraparietal sulcus | 30, -61, 39 | FPN | Fair et al., 2009 |
| L Precun | Left precuneus | -9, -72, 37 | FPN | Fair et al., 2009 |
| R Precun | Right precuneus | 10, -69, 39 | FPN | Fair et al., 2009 |
| L aPFC | Left anterior prefrontal cortex | -28, 51, 1 | CON | Fair et al., 2009 |
| R aPFC | Right anterior prefrontal cortex | 27, 50, 23 | CON | Fair et al., 2009 |
| L aI | Left anterior insula | -35, 14, 5 | CON | Fair et al., 2009 |
| R aI | Right anterior insula | 36, 16, 4 | CON | Fair et al., 2009 |
| dACC | Dorsal anterior cingulate cortex | -1, 10, 46 | CON | Fair et al., 2009 |
| L aTh | Left anterior Thalamus | -12, -15, 7 | CON | Fair et al., 2009 |
| R aTh | Right anterior Thalamus | 10, -15, 8 | CON | Fair et al., 2009 |
| amPFC | Anterior medial prefrontal cortex | 1, 54, 21 | DMN | Fair et al., 2009 |
| vmPFC | Ventromedial prefrontal cortex | -3, 29, -2 | DMN | Fair et al., 2009 |
| L SFC | Left superior frontal cortex | -14, 38, 52 | DMN | Fair et al., 2009 |
| R SFC | Right superior frontal cortex | 17, 37, 52 | DMN | Fair et al., 2009 |
| L ITL | Left inferior temporal lobe | -61, -33, -15 | DMN | Fair et al., 2009 |
| R ITL | Right inferior temporal lobe | 65, -17, -15 | DMN | Fair et al., 2009 |
| L Para(DMN) | Left parahippocampus | -22, -26, -16 | DMN | Fair et al., 2009 |
| R Para(DMN) | Right parahippocampus | 25, -26, -14 | DMN | Fair et al., 2009 |
| pCC | Posterior cingulate cortex | -2, -36, 37 | DMN | Fair et al., 2009 |
| L IPC | Left lateral parietal cortex | -47, -67, 36 | DMN | Fair et al., 2009 |
| R IPC | Right lateral parietal cortex | 53, -67, 36 | DMN | Fair et al., 2009 |
| retsplen | Retro Splenial | 3, 51, 8 | DMN | Fair et al., 2009 |
| supCC | Supragenual cingulate | 0, 34, 30 | EN | Stein et al., 2007 |
| subCC | Subgenual cingulate | 0, 15, -14 | EN | Stein et al., 2007 |
| L OFC | Left orbitofrontal cortex | -46, 31, -9 | EN | Stein et al., 2007 |
| R OFC | Right orbitofrontal cortex | 46, 31, -9 | EN | *Mirrored* |
| L Para(EN) | Left parahippocampus gyrus | -26, -19, -14 | EN | Stein et al., 2007 |
| R Para(EN) | Right parahippocampus gyrus | 26, -19 -14 | EN | *Mirrored* |
| L IPFC | Left lateral prefrontal cortex | -56, 26, 25 | EN | Stein et al., 2007 |
| R IPFC | Right lateral prefrontal cortex | 56, 26, 25 | EN | *Mirrored* |
| L Amyg | Left amygdala | -26, 0, -20 | EN | Stein et al., 2007 |
| R Amyg | Right amygdala | 26, 0, -20 | EN | *Mirrored* |
| pCC | Posterior cingulate cortex | 0, -33, 38 | EN | Stein et al., 2007 |
| L vaI | Left ventral anterior insula | -33, 13, -7 | EN | Deen et al., 2011 |
| R vaI | Right ventral anterior insula | 32, 10, -6 | EN | Deen et al., 2011 |
| mPMC | Mesial prefrontal cortex | -3 -5 60 | MN | Mayka et al., 2006 |
| pSMA | pre-supplementary motor area | -4 3 59 | MN | Mayka et al., 2006 |
| SMA | Supplementary motor area | -3 -11 61 | MN | Mayka et al., 2006 |
| L lPMC | Left lateral premotor cortex | -26 -10 62 | MN | Mayka et al., 2006 |
| R lPMC | Right lateral premotor cortex | 25, -11, 61 | MN | *Mirrored* |
| L dlPMC | Left dorsolateral premotor cortex | -30 -8 64 | MN | Mayka et al., 2006 |
| R dlPMC | Right dorsolateral premotor cortex | 29 -9 64 | MN | *Mirrored* |
| L vlPMC | Left ventral lateral premotor cortex | -52 7 22 | MN | Mayka et al., 2006 |
| R vlPMC | Right ventral lateral premotor cortex | 52 6 21 | MN | *Mirrored* |
| L SMC | Left sensorimotor cortex | -39 -24 59 | MN | Mayka et al., 2006 |
| R SMC | Right sensorimotor cortex | 38 -25 59 | MN | *Mirrored* |
| L M1 | Left primary motor cortex | -37 -25 64 | MN | Mayka et al., 2006 |
| R M1 | Right primary motor cortex | 36 -21 58 | MN | *Mirrored* |
| L S1 | Left primary sensorimotor cortex | -40 -27 54 | MN | Mayka et al., 2006 |
| R S1 | Right primary sensorimotor cortex | 39 -28 54 | MN | *Mirrored* |
| R Ling | Lingual_R | 18, -47, -10 | VN | Cassady et al., 2019 |
| R Ling | Lingual_R | 20, -66, 2 | VN | Cassady et al., 2019 |
| R Ling | Lingual_R | 20, -86, -2 | VN | Cassady et al., 2019 |
| L Ling | Lingual_L | -15, -72, -8 | VN | Cassady et al., 2019 |
| L Ling | Lingual_L | -16, -52, -1 | VN | Cassady et al., 2019 |
| R Cal | Calcarine_R | 8, -72, 11 | VN | Cassady et al., 2019 |
| R Cal | Calcarine_R | 6, -81, 6 | VN | Cassady et al., 2019 |
| L Cal | Calcarine_L | -18, -68, 5 | VN | Cassady et al., 2019 |
| L Cal | Calcarine_L | -8, -81, 7 | VN | Cassady et al., 2019 |
| R Cun | Cuneus_R | 15, -77, 31 | VN | Cassady et al., 2019 |
| R Cun | Cuneus_R | 6, -72, 24 | VN | Cassady et al., 2019 |
| L Cun | Cuneus_L | -16, -77, 34 | VN | Cassady et al., 2019 |
| L Cun | Cuneus_L | -3, -81, 21 | VN | Cassady et al., 2019 |
| R Fus | Fusiform_R | 27, -59, -9 | VN | Cassady et al., 2019 |
| L Fus | Fusiform_L | -33, -79, -13 | VN | Cassady et al., 2019 |
| R SupOcc | Occipital_Sup_R | 15, -87, 37 | VN | Cassady et al., 2019 |
| R SupOcc | Occipital_Sup_R | 24, -87, 24 | VN | Cassady et al., 2019 |
| L SupOcc | Occipital_Sup_L | -14, -91, 31 | VN | Cassady et al., 2019 |
| R MidOcc | Occipital_Mid_R | 40, -72, 14 | VN | Cassady et al., 2019 |
| R MidOcc | Occipital_Mid_R | 29, -77, 25 | VN | Cassady et al., 2019 |
| R MidOcc | Occipital_Mid_R | 37, -81, 1 | VN | Cassady et al., 2019 |
| R MidOcc | Occipital_Mid_R | 37, -84, 13 | VN | Cassady et al., 2019 |
| L MidOcc | Occipital_Mid_L | -40, -88, -6 | VN | Cassady et al., 2019 |
| L MidOcc | Occipital_Mid_L | -28, -79, 19 | VN | Cassady et al., 2019 |
| L MidOcc | Occipital_Mid_L | -24, -91, 19 | VN | Cassady et al., 2019 |
| L MidOcc | Occipital_Mid_L | -42, -74, 0 | VN | Cassady et al., 2019 |
| L MidOcc | Occipital_Mid_L | -26, -90, 3 | VN | Cassady et al., 2019 |
| R InfOcc | Occipital_Inf_R | 43, -78, -12 | VN | Cassady et al., 2019 |
| L InfOcc | Occipital_Inf_L | -47, -76, -10 | VN | Cassady et al., 2019 |
| R InfTemp | Temporal_Inf_R | 42, -66, -8 | VN | Cassady et al., 2019 |
| L InfTemp | Temporal_Inf_L | -42, -76, -10 | VN | *Mirrored* |
| R Hes | Heschl_R | 32, -26, 13 | AN | Cassady et al., 2019 |
| L Hes | Heschl_L | -30, -27, 12 | AN | Cassady et al., 2019 |
| L Hes | Heschl_L | -55, -9, 12 | AN | Cassady et al., 2019 |
| R Sup Temp | Temporal_Sup_R | 65, -33, 20 | AN | Cassady et al., 2019 |
| R Sup Temp | Temporal_Sup_R | 58, -16, 7 | AN | Cassady et al., 2019 |
| L Sup Temp | Temporal_Sup_L | -60, -25, 14 | AN | Cassady et al., 2019 |
| L Sup Temp | Temporal_Sup_L | -49, -26, 5 | AN | Cassady et al., 2019 |
| R RolOp | Rolandic_Oper_R | 43, -23, 20 | AN | Cassady et al., 2019 |
| R RolOp | Rolandic_Oper_R | 56, -5, 13 | AN | Cassady et al., 2019 |
| L RolOp | Rolandic_Oper_L | -38, -33, 17 | AN | Cassady et al., 2019 |
| R SupMar | SupraMarginal_R | 59, -17, 29 | AN | Cassady et al., 2019 |
| L SupMar | SupraMarginal_L | -50, -34, 26 | AN | Cassady et al., 2019 |
| L SupMar | SupraMarginal_L | -53, -22, 23 | AN | Cassady et al., 2019 |

**Supplemental Table 2.** All lateralized ROIs without a contralateral match were mirrored in the x direction. The following statistics are partial Pearson’s correlation coefficients measuring the association between the GE of CB-BG networks and each affected network using only original ROIs (no mirrored)

|  | EN | MN | VN |
| --- | --- | --- | --- |
| CCB-BG | .16* | .18** | .10 |
| MCB-BG | .03 | .12 | .28*** |

*Note: Analyses were assessed using FDR correction. \*\*\* $p < .001$ \*\* $p < .01$ \* $p < .05$*

**Supplemental Table 3.** Pearson Correlation Coefficients (*r*) showing the relationship between each cortical network’s global efficiency.

|  | FPN | CON | DMN | EN | MN | VN | AN |
| --- | --- | --- | --- | --- | --- | --- | --- |
| FPN | 1 | .21** | .19** | .13 | .14* | .17* | .25*** |
| CON | .21** | 1 | .27*** | .20** | .30*** | .28*** | .35*** |
| DMN | .19** | .27*** | 1 | .25*** | .31*** | .27*** | .25*** |
| EN | .13 | .20** | .25*** | 1 | .24*** | .20** | .25*** |
| MN | .14* | .30*** | .31*** | .24*** | 1 | .39*** | .39*** |
| VN | .17* | .28*** | .27*** | .20** | .39*** | 1 | .40*** |
| AN | .25*** | .35*** | .25*** | .25*** | .39*** | .40*** | 1 |

*Note: Analyses were assessed using FDR correction. \*\*\* $p < .001$ \*\* $p < .01$ \* $p < .05$*

